# In-vitro Modeling of Intravenous Drug Precipitation by the Optical Spatial Precipitation Analyzer (OSPREY)

**DOI:** 10.1101/2023.03.02.530827

**Authors:** Andrew J. Radosevich, Ruth L. Martin, Wayne R. Buck, Lauren Hicks, Amanda Wilsey, Jeffrey Y Pan

## Abstract

Intravenous (IV) administration of poorly water-soluble small molecule therapeutics can lead to precipitation during mixing with blood. This can limit characterization of pharmacological and safety endpoints in preclinical models. Most often, tests of kinetic and thermodynamic solubility are used to optimize the formulation for solubility prior to infusion in animals, but these do not capture the dynamic precipitation processes that take place during *in-vivo* administration. To better capture the fluid dynamic processes that occur during IV administration, we developed the Optical Spatial Precipitation AnalYzer (OSPREY) as a method to quantify the amount and size of compound precipitates in whole blood using a flow-through system that mimics IV administration. Here, we describe the OSPREY device and its underlying imaging processing methods. We then validate the ability to accurately segment particles according to their size using monodisperse suspensions of microspheres (diameter 50 to 425 microns). Next, we use a tool compound, ABT-737, to study the effects of compound concentration, vessel flow rate, compound infusion rate and vessel diameter on precipitation. Finally, we use the physiological diameter and flow rate of rat femoral vein and dog saphenous vein to demonstrate the potential of OSPREY to model *in-vivo* precipitation in a controlled, dynamic *in-vitro* assay.

**Highlights:**

- Prospective small molecule therapeutics are often solubility challenged when injected into whole blood at elevated concentrations for toxicology studies.
- Improved *in-vitro* solubility measurements in a flowing system are needed to better understand *in-vivo* intravenous precipitation
- OSPREY is a novel *in-vitro* flow-through system that quantifies solubility in whole blood

**Graphical Abstract:** 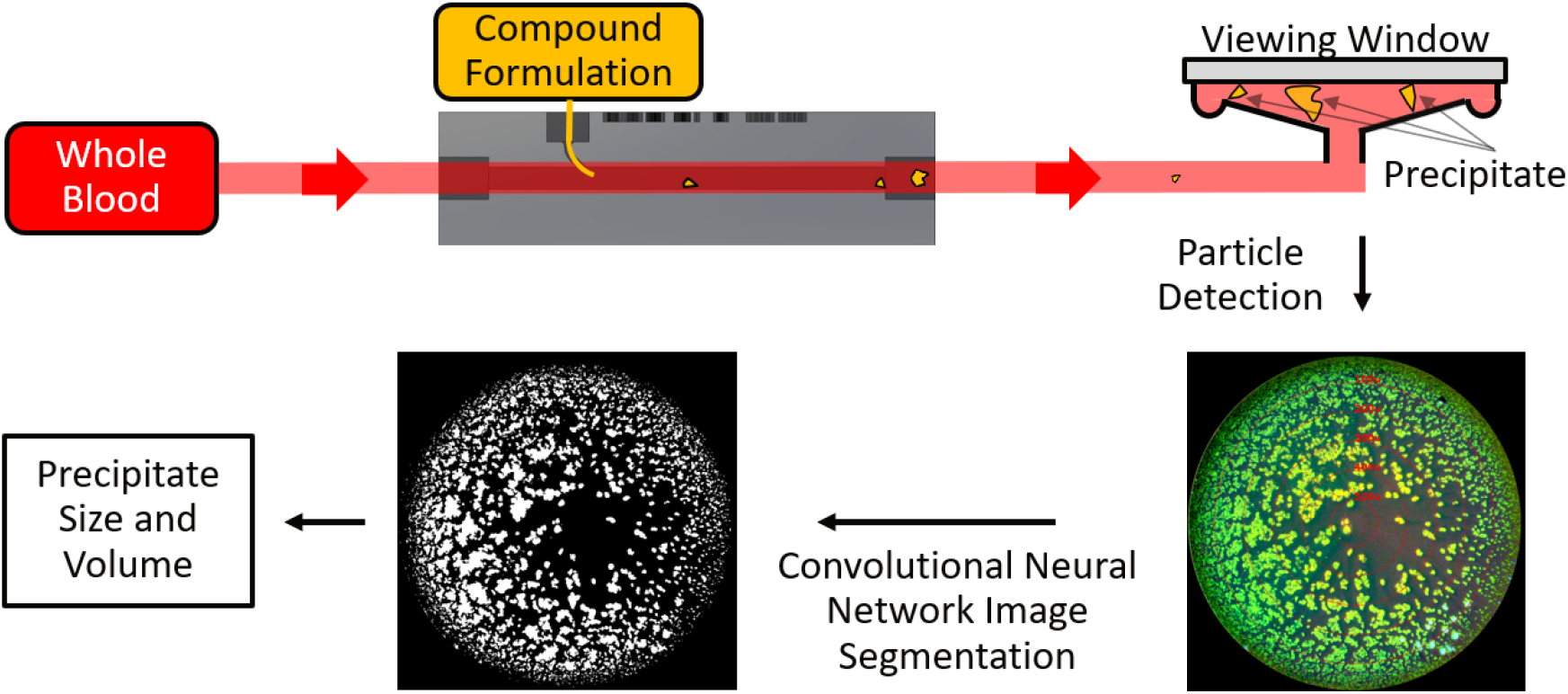

## 1 Introduction

Changes in medicinal chemistry approaches used in the pharma industry, including the adoption of combinatorial chemistry and high throughput screening, have led to reduced solubility of compounds during hit generation(Stegemann et al., 2007). As a result, enhancement of drug-like properties, including solubility, is becoming a more important aspect of lead optimization during early drug discovery(Das et al., 2022). A particular challenge is formulating aqueous solubility challenged compounds in such a way that they can be studied in preclinical models via IV administration without forming precipitate as they enter the blood stream. To overcome such issues, a number of comple formulation approaches have been described, such as prodrugs and lipid nanoparticles, which can enable delivery of poorly soluble compounds(Shi et al., 2009). However, preclinical exploration of safety often relies on early unoptimized leads, or administration of compounds by parenteral routes not indicated in the clinic(Koshman et al., 2018; Kym et al., 2006). As a result, development costs associated with specialized IV formulations prevent their routine application during early discovery safety evaluation.

Although strongly hydrophilic vehicles, such as PEG-400, can solubilize compounds for IV administration, these formulations can cause IV precipitation resulting in adverse effects (Yalkowsky et al., 1983) or impaired pharmacokinetics. The best documented risk of IV infusion is the introduction of parenteral nutrition mixtures containing unintended calcium phosphate crystals or the formulation of calcium phosphate crystals upon the warming of these solutions in the blood(McKinnon, 1996). Hill and colleagues reported the unfortunate death of two young women who received IV parenteral nutrition containing invisible calcium phosphate precipitate resulting in respiratory failure and death, with embolization of pulmonary microvasculature by calcium phosphate crystals confirmed postmortem(Hill et al., 1996). Similarly, high-resolution CT was able to document the pulmonary localization of radiopaque crystals from a different parenteral nutrition mixture which biopsy revealed were emboli within pulmonary microvessels(Reedy et al., 1999). These literature reports are consistent with our own observations that IV administration of compounds which precipitate intravenously result in pulmonary embolization that can cause cardiovascular collapse in an anesthetized dog model. As a result, the desired measurement of pharmacological effects of the drug can become obscured by precipitation driven by the physicochemical properties of the compound. To avoid such uninformative studies, we sought to understand the limits of concentration and infusion rate for infusion into our preclinical models to achieve the highest possible IV compound exposure without experiencing adverse effects due to precipitation.

Evaluation of the propensity of a compound formulation to precipitate when mixed in an 1 aqueous environment can be performed by assaying sample aliquots in an agitated vessel system. However, such assay systems fail to capture the complex fluidic dynamics that occur in a flowing *in-vivo* system. To assess the flowing dynamic precipitation of small molecule formulations in an *in-vitro* setting, Evans *et al* developed a system that injected compound formulations and flowed blood surrogate (Isotonic Sorensen’s phosphate buffer) through a commercially available 4 mm diameter round quartz tube using an IV Y-Site^2^. Sun *et al*, developed a similar flow-through system using two quartz plates separated by a spacer, with precipitation and concentration measured using UV-vis imaging and a UV ^3^spectrophotometer. We sought to build upon these models to enable the use of whole blood, modulate flow cell design to match multiple relevant physiological structures, characterize precipitate size in addition to total precipitate volume, and integrate programmatic controls to automate relevant system parameters including compound concentration, flow rates, and quantitative analysis.

We developed the Optical Spatial PREcipitation analYzer (OSPREY) as a method to automate the quantitation of precipitation events in flowing whole blood. OSPREY consists of a three-syringe system to model blood flow and compound infusion, a 3D printed injection block (IB) that enables us to explore different infusion geometries, a conical particle detector (CPD) to quantify precipitate sizing through whole blood, and an image processing software that uses a convolutional neural network algorithm to automatically quantify compound precipitates.

In this work, we begin with an introduction to the OSPREY hardware and image analysis that provides automated particle detection using U-NET convolutional neural network (CNN) image segmentation(Ronneberger et al., 2015). We then validate OSPREY’s ability to accurately quantify particles of different size using monodisperse suspensions of microspheres from 50 to 425 μm in diameter. Next, we demonstrate the versatility of our approach using a tool compound (ABT-737) to study the effects of compound concentration, vessel flow rate Q_v_, infusion flow rate Q_i_, and vessel diameter D_v_ on the amount and size of precipitates formed. We finish by simulating the physiological flow rates and diameters for the *in-vivo* IV administration of ABT-737 into a dog saphenous vein and rat femoral vein.

## Materials and Methods

### 2.1 OSPREY Instrumentation

The OSPREY instrument consists of a CPD (Fig. 1B), a transparent 3D printed compound IB (Fig. 1E), and a 3-syringe pump system to model steady IV blood flow and compound injection flow. Blood or an appropriate surrogate liquid is aspirated into the first glass syringe and subsequently dispensed at a constant flow rate across the IB and into the CPD. The second and third glass syringes introduce vehicle-diluted compound solution into the main liquid line via a catheter inserted into the IB. Compound precipitates that form in the system travel downstream into the CPD. Here, particles suspended in the model blood flow out from the center of the device and become trapped at a radius that depends on their size, with larger particles appearing towards the center and smaller particles towards the edge (Fig. 1C – with medium size particle depicted). A color scientific camera (Thorlabs, Zelux CS165CU) records video of the compound injection site and a second color scientific camera (Thorlabs, Kiralux CS505CU) records compound precipitate distributions trapped in the CPD. The CPD is evenly illuminated with an oblique angle white light LED (Thorlabs, MWWHLP2) and enclosed within a black box to eliminate background light. 3D computer-aid design (CAD) models for the CPD and IB are included in the Supplementary Information.

**Fig. 1.**
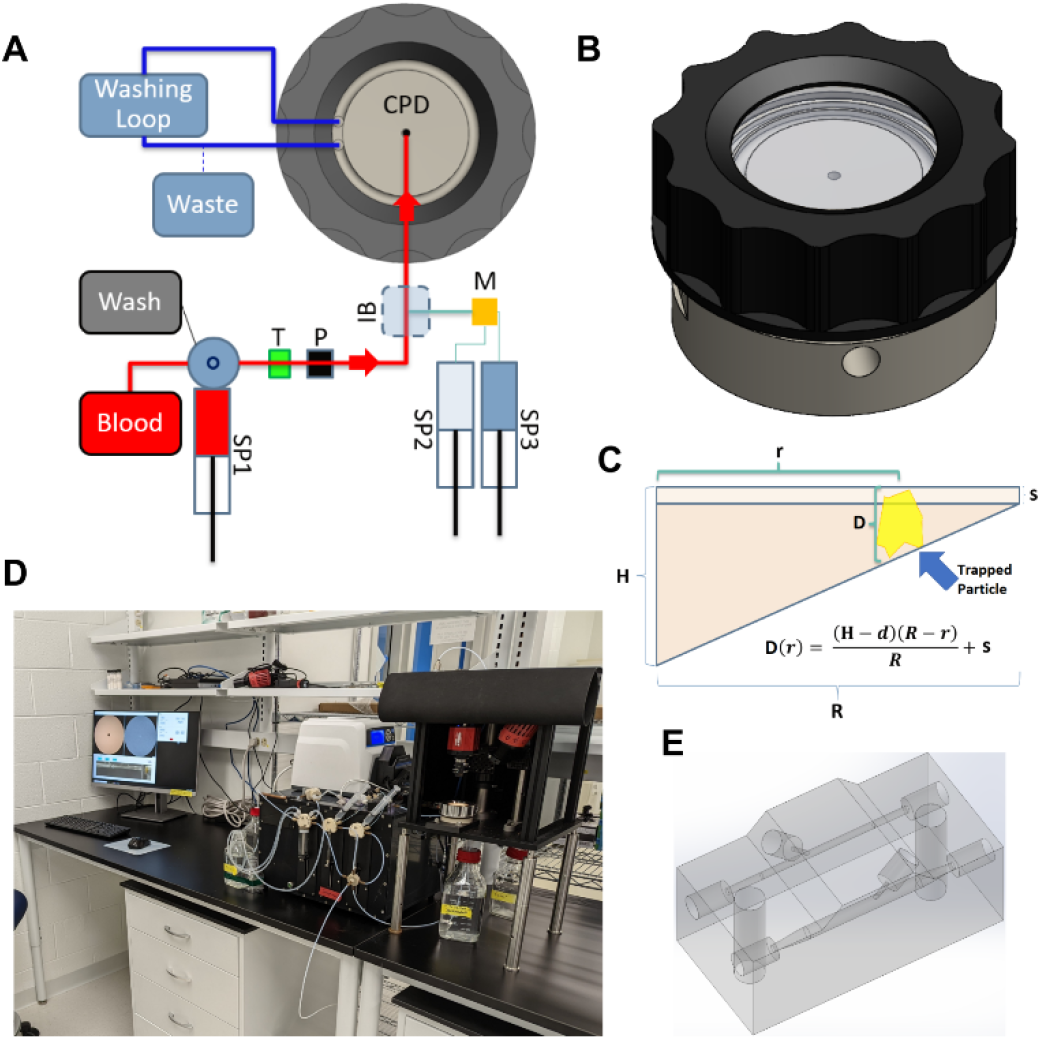
Hardware components of the OSPREY instrument. A) Device schematic with syringe pumps SP, temperature sensor T, pressure sensor P, injection block IB, and mixer M. B) 3D rendering of the conical particle detector CPD. C) Schematic showing the parameters used to calculate the precipitate diameter D based on its trapping radius where H is the distance from the face of the quartz window to the surface of the conical section at the central inlet port, s is the distance from the quartz window to the edge of the conical section, R is the outer radius of the conical surface, and r is the measured distance away the center in which a particle becomes trapped. A trapped particle at distance r is shown in yellow. D) Picture of the full OSPREY instrument sitting on a lab bench. E) 3D rendering of the IB.

The CPD consists of a lower conical recess cut into PEEK, a flat Quartz disc viewing window (McMaster-Carr, 1357T32), a silicone O-ring (McMaster-Carr, 1173N032) along the outer rim of the detector, and a threaded aluminum cap. Model blood flows up into the conical section through a 0.102” (∼2.59 mm) diameter inlet orifice and travels out radially until spilling over into a collection moat outside of the 1.456” (∼37 mm) diameter outer ring of the conical section. The maximum vertical separation between the conical recess and Quartz viewing window is 0.021” (∼533 μm; H in Fig. 1C) at the edge of the inlet orifice and 10 μm at the outer ring of the conical section (s in Fig. 1C). The vertical separation between the conical surface and flat viewing window varies linearly between H and s over a horizontal distance R. As a result, the particle diameter can be determined from its trapping radius r according to:

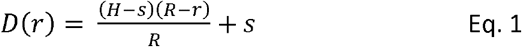

In other words, the effective size of any compound precipitate can be determined by identifying the radius away from the center at which it becomes trapped. As a result, the CPD can be used to quantify the total volume and size distribution of compound particles that are formed in the OSPREY device.

The minimum quantified particle size is set by the smallest separation between the conical surface and flat viewing window of the CPD (e.g., s = 10 μm). Particles larger than this size are trapped in the CPD and quantified, whereas smaller particles flow unimpeded through the system. We chose 10 μm since this is roughly the diameter of a capillary and represents the lower end of particle size that would cause vascular blockages *in-vivo*.

A major advantage of the CPD is the ability to assess compound formulations infused into whole blood. Whole blood is largely opaque to visible light over distances larger than a few hundred microns. As a result, it is not possible to view inside a vessel on the order of a few millimeters using optical imaging. OSPREY overcomes this issue by using the CPD to trap particles against a quartz window for analysis, thereby minimizing the optical pathlength through which the probing light passes through whole blood.

The IB is 3D printed in Accura ClearVue by Quickparts. The model blood vessel has a circular cross section that runs straight across the 1.8” length of the piece and is terminated at either end by ¼-28 threaded holes. The lumen is treated with a clear coat to minimize the inherent surface roughness of the 3D printing process. Multiple IBs were made with model blood vessel diameters D_v_ = 1.6 mm, 2.03 mm, 2.45 mm, and 2.9 mm. A 1 mm diameter injection hole enters the model blood vessel at a 45° angle of incidence. An injection catheter tube (McMaster-Carr 5335K9, 305 μm inner diameter, 762 μm outer diameter, ∼33 mm Length, ∼2.4 μL inner volume) is inserted through the injection hole and is sealed by a fluidic fitting (IDEX P-259). The outlet of the injection catheter is positioned at the center of the model blood vessel.

The 3-syringe pump (Cavro, XLP 6000) flow system consists of a 25 mL syringe pump SP1 that provides model blood flow and two 100 μL syringe pumps SP2 and SP3 whose flow streams are combined in a low-volume mixing T (IDEX U-466S, 2.2 μL swept volume) to introduce compound at a user specified concentration into the model blood vessel. Since venous blood flow is not pulsatile, a constant flow rate is provided by SP1 to mimic *in-vivo* venous blood flow. The infusion of compound from SP2 and SP3 is delayed by 1 second after the start of SP1 to allow the model blood to reach a steady state. In-line sensors monitor the model blood temperature (Pendotech, TEMPS-N-012) and pressure (Pendotech, PREPS-N-000). The first of the two 100 μL syringes contains compound at the highest concentration under analysis, which is typically the solubility limit of the given compound in the corresponding vehicle. The second of the two 100 μL syringes contains diluent. By controlling the relative flow rate between the two syringes through control software, users can dynamically specify the final concentration of compound that is introduced into the model vessel flow stream.

The flow path volume between the compound infusion site and the inlet port of the CPD is fixed at ∼0.5 mL. Therefore, to a first approximation, the timescale on which we detect precipitates is ∼0.5 / Q_v_ minutes. Using the range of flow rates presented in this work gives residence times between 0.46 - 7.5 seconds.

By changing the syringe diameters on SP1, SP2, and SP3 different volumes and flow rate ranges can be accessed. Here, we choose syringe diameters that enable flow rates representative of those found in preclinical animal studies. For SP1, the nominal flow rate ranges from 0.15625 mL/min to 1250 mL/min for water. For SP2 and SP3, the nominal flow rate ranges from 6.25 × 10^4^ mL/min to 5 mL/min for water. Since whole blood and compound solutions are more viscous that water, the actual upper flow rate that is possible without stalling the syringe motor is likely lower. Therefore, when changing to a new model system, some experimentation to choose the optimal syringe barrel diameter may be needed.

All surfaces of the flow path that come in contact with blood or compound solution were chosen for their bio- and chemical compatibility. This includes glass syringes with PTFE selector values, platinum-cured silicone tubing for aspirating blood, a biocompatible clear coat on the lumen of the IB, and a PEEK and glass CPD. An automated washing routine performed between experiments flows 0.1% Triton X detergent through the flow path in the reverse direction. This cleans the surfaces of the flow path and removes previously trapped particles. To ensure proper long-term function, each instrument component is easily dissembled and manually cleaned as needed throughout the experiment day.

### 2.2 Image Processing

A minimum of 10 images are collected over the time course of the compound injection. To normalize for debris from prior experiments, machining artifacts on the detector surface, and inhomogeneities in illumination, images are presented for analysis after subtracting a clean reference image (Figs. 2A and 2B) from each subsequent frame where precipitate is collected in the detector (Figs. 2C and 2D). Such differential images (Figs. 2E and 2F) optimize the contrast for quantifying newly formed particle aggregates. This approach works similarly well for both transparent fluids such as plasma (Figs. 2A, 2C, and 2E) and whole blood (Figs. 2B, 2D, and 2F). The differential image was calculated for each channel of a color image separately and so retains some of the spectral information from the sample that makes it easier to distinguish compound precipitate from other potential contamination (e.g., bubbles, fibers, dust, etc.). A 25x speed video time course of the differential image over 11 experiments with escalating concentration (0 to 50 mg/mL) for both plasma and whole blood are included in the Appendix.

**Fig. 2.**
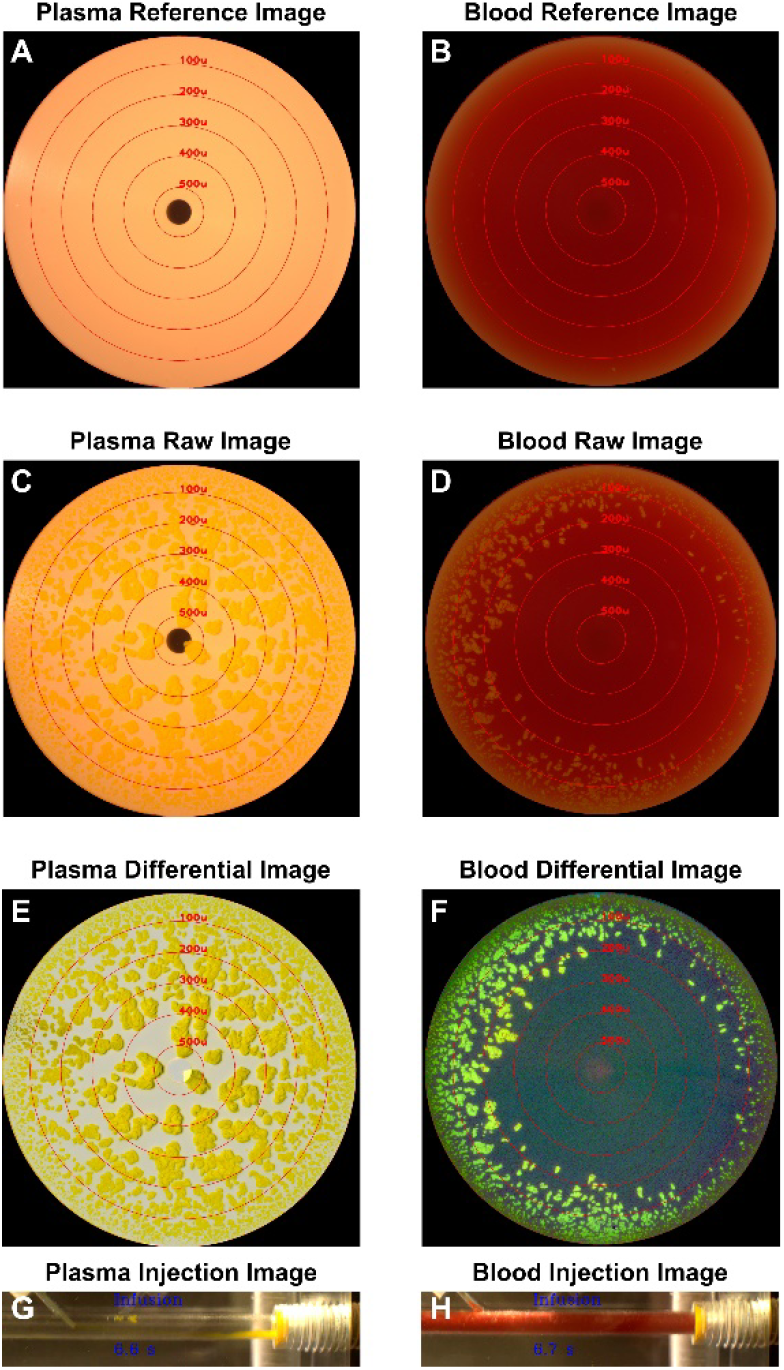
Images of the CPD and IB during the injection of 50 mg/mL ABT-737 at 100 μL/min into dog plasma (left column) and whole bovine blood (right column) at 10 mL/min flow rate. Clean reference image of the CPD in plasma A) and whole blood B). Precipitate filled image of the CPD at the end of compound infusion for plasma C) and whole blood D). Differential image that subtracts the reference image from the precipitate filled image of the CPD in plasma E) and whole blood F). IB images during compound infusion for plasma G) and whole blood H).

Video of the IB is recorded throughout the experiment with representative still frames shown in Figs. 2G and 2H for plasma and whole blood, respectively. The corresponding real-time videos are included in the Appendix. In plasma, it is possible to record precipitate forming near the injection site, whereas whole blood is opaque to visible light so precipitate formation cannot be viewed. Despite this advantage to acquiring measurements in a transparent whole blood surrogate, we found that the size and amount of precipitation differs between plasma and whole blood (Figs. 2E and 2F). As a result, we use whole blood for the remaining studies of ABT-737 in this paper to provide the results most comparable to *in-vivo* conditions.

### 2.3 Automated precipitate detection using CNN segmentation

To automatically identify compound precipitate boundaries in the differential images, we trained a CNN image segmentation algorithm using the U-Net architecture.(Ronneberger et al., 2015) The network architecture and full Python processing code using Python 3.7.4, Keras 2.2.5, and Tensorflow 1.14.0 are included in the Supplementary Information. To train the CNN, we first manually segmented tens of images using the ‘Masks from ROIs’ plugin in FIJI ImageJ or used an Otsu’s method thresholding in MATLAB to generate binary ground truth images(Schindelin et al., 2012; Schneider et al., 2012). The full resolution differential (Figs. 3A and 3D) and ground truth (Figs. 3B and 3E) images were then split into 16 separate 512×512 pixel images and trained to find the optimal network weights. The fitting was performed with various hyperparameters until the predicted masks (Figs. 3C and 3F) for a reserved testing set of images were assessed to be qualitatively similar to their corresponding ground truth images. A full 2048 × 2048-pixel image is segmented in under 2 seconds on an Intel Core i9-9880H CPU. Figure 3 shows a representative demonstration of the ability of the developed CNN to predict precipitate areas with good qualitative agreement with the ground truth image. For this image, 94.2% of the yellow pixels in the ground truth image were accurately predicted by the CNN. Further examples in the reserved testing set are included in the SI and show similarly good agreement.

**Fig. 3.**
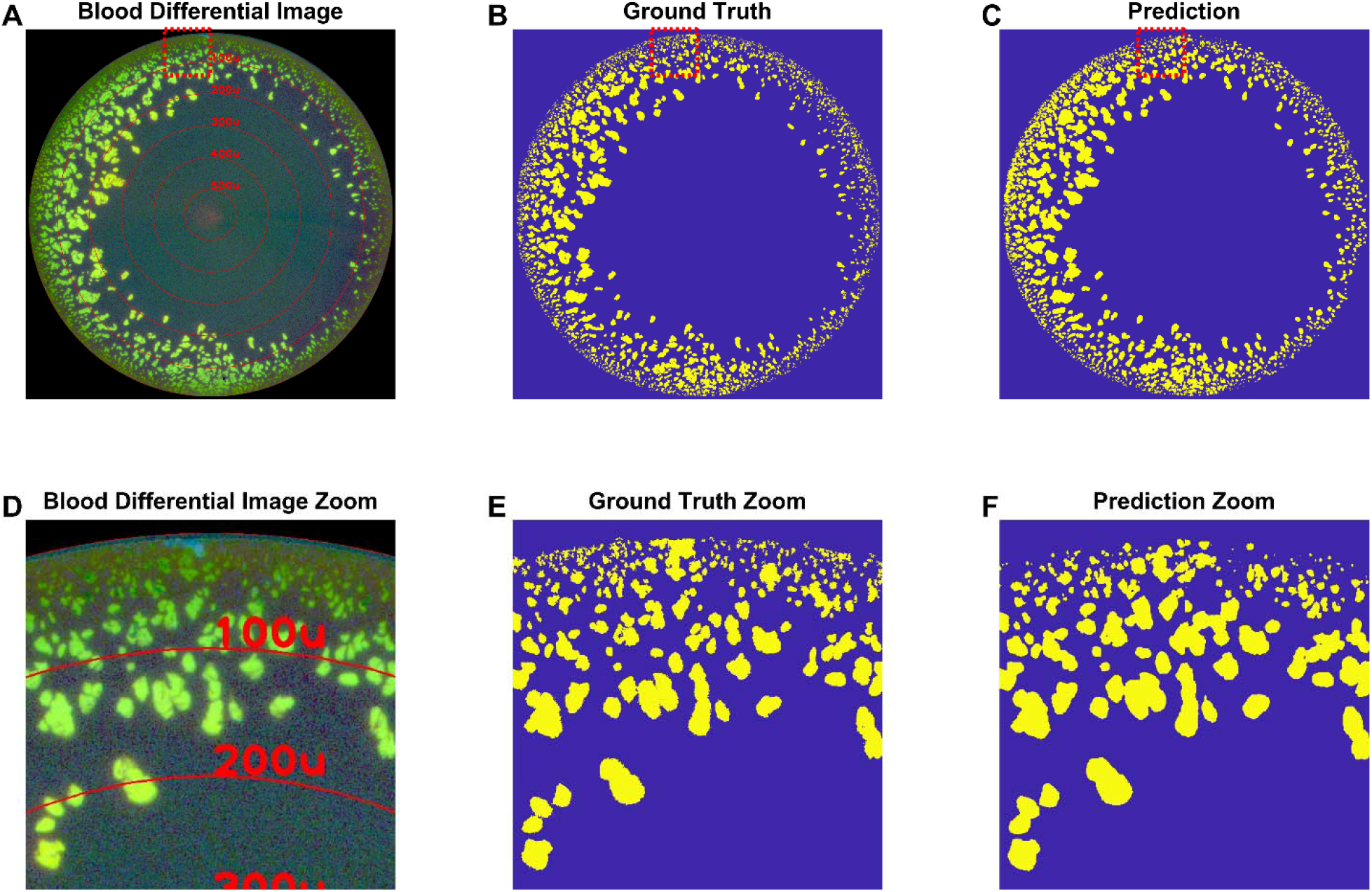
Demonstration of automatic precipitate area detection of ABT-737 particles in whole blood using the UNET architecture convolutional neural network (CNN). A) Differential image of precipitate formed with 50 mg/mL concentration, 100 μL/min infusion rate, and 10 mL/min blood flow rate. B) Ground truth (manual) segmentation image showing areas with precipitate in yellow. C) Predicted precipitate areas after CNN analysis. The differential and ground truth image were not included in the training data for the CNN, so the prediction was naïve to this image set. Panels D), E), and F) show the zoomed in images of the area highlighted by the red dashed square in panels A), B), and C).

Using a CNN to perform image processing has several benefits. First, it limits operator bias by automatically applying the same processing algorithm across a dataset. Second, it overcomes the sensitivity of traditional intensity thresholding approaches to unwanted artifacts (e.g., fibers, blood clumps, etc.). Third, new compound that may exhibit different color, shape, or morphology can be quickly trained by supplying the algorithm with additional curated datasets. Finally, it allows for the future possibility to both segment and classify particles into different categories.

### 2.4 Quantification of total particle volume and characteristic size

To quantify the amount and size of precipitate trapped in the CPD, we developed metrics for the total particle volume *V*_*T*_ and the characteristic particle size Lc from first principles. The vertical separation D between the CPD surface and flat viewing window varies radially about the center of the image according to Eq 1. To estimate the volume *V* of precipitate that could be trapped at each pixel location, we approximate each pixel’s volume as a rectangular prism with height *D*, width *w*, and length *l*:

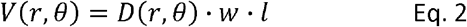

where *w* and *l* are constant across all pixels and are calculated empirically using the diameter of the CPD as a reference length scale. Converting to cartesian coordinates and applying a mask with the particle locations predicted by the CNN (e.g., Fig. 3C), the radial particle volume distribution *V*(*r*) is:

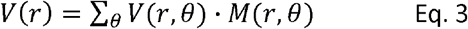

The distribution of particle volume as a function of particle diameter *V*(*D*) can then be found by transforming from the radial coordinate space of *V*(*r*) into particle diameter space using Eq. 1. *V*_*T*_ is then determined by summing the contribution from all particle diameters:

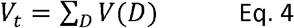

*L*_*c*_ is calculated as the first moment of *V*(*D*):

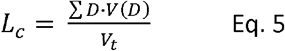

Conceptually, *V*_*t*_ represents the aggregate volume of particles trapped in the CPD and is expressed here in units of μL. *L*_*c*_ can be thought of as the average trapped particle size expressed in units of μm. Therefore, when relatively small particles are formed and trapped in the CPD, *L*_*c*_ will be a smaller value than when larger particles are trapped.

### 2.5 Whole blood and plasma preparation

Whole bovine blood (Innovative Research, IGBOWBK2E 500ML) or defibrinated horse blood (Hardy Diagnostics, DHB500) was filtered with a 100 μm nylon cell strainer (Corning, 431752) prior to analysis and gently stirred at room temperature to keep the blood evenly mixed. Dog plasma was prepared from whole blood collected from acutely anesthetized Beagle dogs just prior to euthanasia, using a single collection blood bag, stored at 4°C, and processed within 24 hours of collection. Blood was centrifuged for 3 hours at 20°C, 2000 x g in a Beckman Coulter, Allegra X-12 centrifuge. The resulting supernatant was centrifuged for another 3 hours at the previous settings to further reduce red blood cell contamination. The resulting plasma was stored at -80°C until use. All procedures were approved by AbbVie’s Institutional Animal Care and Use Committee and carried out in American Association for Accreditation of Laboratory Animal Care-accredited facilities.

### 2.6 Monodisperse microsphere standard

Near water density (∼1.00 g/mL) fluorescent green polyethylene microspheres (Cospheric, UVPMS-BG-1.00) of diameter 50 μm, 180 μm, 300 μm, and 425 μm were suspended in water with 0.1% Triton X detergent to prevent the particles from clumping. Microsphere suspensions were mixed with a magnetic stir bar prior to aspiration into SP1 and dispensed through the IB into the CPD at 10 mL/min flow rate.

### 2.7 Small molecule sample and vehicle

ABT-737 is an early BCL-2 inhibitor(Oltersdorf et al., 2005) for which we observed in vivo precipitation in animal species after IV infusion of PEG-400-based formulations (data not shown). The compound concentrations achievable in plasma, despite poor aqueous solubility, are due to high protein binding to albumin(Vogler et al., 2010). ABT-737 is a zwitterionic compound at neutral pH that rapidly forms a yellow amorphous precipitate when PEG-400 formulations are diluted into aqueous solvents or biofluids. Therefore, it was selected as a tool compound for determining instrument parameters which alter precipitation kinetics.

To prepare the high concentration syringe for SP2, 200 μL of warmed Dimethylacetamide was added to 50 mg of ABT-737 to fully dissolve the compound. An additional 800 μL of PEG-400 was added and the sample was vortexed. The solution was aspirated into a 1 mL plastic syringe and loaded onto OSPREY.

### 2.8 Physiological model systems

Dog saphenous vein measurements of Q_v_ = 65 mL/min and D_v_ = 2.9 mm were acquired by Doppler ultrasound (data not shown). Compound was infused with Q_i_ = 100 μL/min to model a 10 kg dog receiving an infusion at a rate of 0.01 ml/kg/min(Koshman et al., 2018). Rat femoral vein D_v_ = 1.6 mm was estimated from *ex-vivo* vessel samples and Q_v_ = 2.5 mL/min was obtained by scaling down the flow velocities found in human(Abraham et al., 1994). Compound was infused with Q_i_ = 75 μL/min to model a 350 g rat receiving a 1 ml/kg dose volume over a 4-minute slow bolus, which is consistent with current practice of organic formulation infusions(Banfor et al., 2016).

## 3 Results and Discussion

### 3.1 Validation of particle sizing in the conical particle detector

Monodisperse suspensions of microspheres were used to validate the ability of the CPD to accurately quantify particle diameter across the detector’s field of view. The microsphere suspensions were loaded into SP1 and dispensed at 10 mL/min through the full flow-through system where they became trapped in circular ring regions around the central inlet port of the CPD. The size of each microsphere can be determined by measuring the ring radius in which they are trapped and a priori knowledge of the device dimensions as depicted in Fig. 1C. Fig. 4A shows the raw image of trapped microspheres with nominal diameter of 50 μm, 180 μm, 300 μm, and 425 μm. This image was segmented using Otsu’s method and the polar coordinates centroid of each microsphere found. Fig. 4B shows blue dots corresponding to the centroid of each microsphere plotted as a function of diameter and azimuthal angle around the central inlet port. Most of the microspheres are found with the nominal diameter variation expected for these microspheres. Moreover, the trend is relatively flat across the azimuthal angle, indicating that the CPD is machined symmetrically about the central port.

**Fig. 4.**
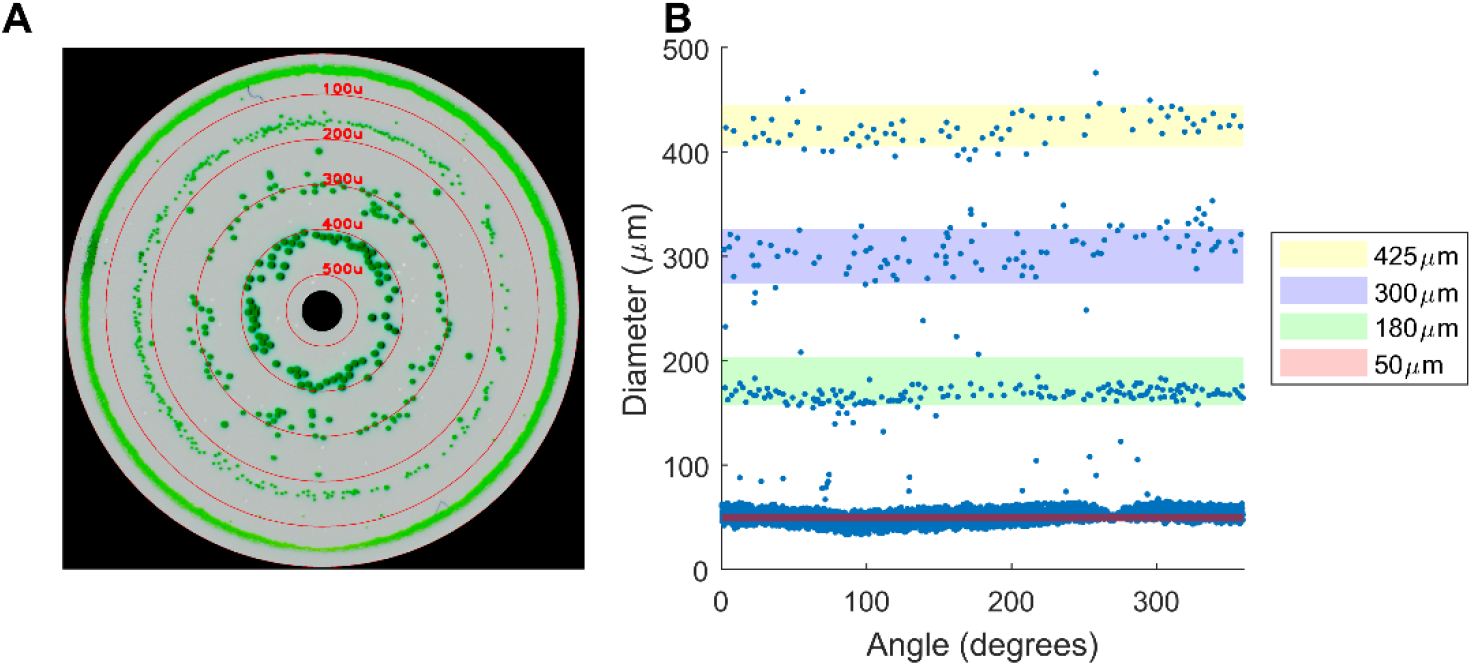
Demonstration of particle sizing capability of the CPD. A) Detector image showing 4 concentric rings corresponding to green fluorescent microspheres of diameter 50 μm, 180 μm, 300 μm, and 425 μm. B) Plot showing the diameter of each particle as a function of azimuthal angle around the detector.

Qualitatively, the lateral size of individual microspheres imaged by the camera is larger the closer they are trapped towards the central inlet port of the CPD (Fig. 4A). A similar effect is seen in actual compound precipitates (e.g., Figs. 2E and 2F). Although in principle the lateral size could also be used to quantify particle sizing, actual compound precipitates are more jagged and overlapping than microspheres. This makes it difficult to categorically determine individual particle boundaries for sizing. As a result, we have found that quantifying the size distribution according to Eq. 3 is a more straightforward and robust method.

For the 50 μm spheres, the range of calculated centroid diameters is broader than the range of sizes (indicated by the red band in Fig. 4B) expected according to the manufacturer’s specifications. This is due in part to more 50 μm spheres being loaded into the CPD than there is physical space to allow for. As a result, spheres can become bunched up at a location that is not representative of their actual size. In other words, particles can become trapped by a downstream blockage rather than being wedged between the conical section and quartz window if the particle concentration becomes too high. Because of this potential effect, it is important to not overload the detector, otherwise the ability to accurately quantify size is diminished. If overloading is suspected in a particular experiment, we can simply decrease the infused volume to confirm the results are robust.

### 3.2 Effect of Varying Compound Concentration, Vessel Flow Rate, Injection Flow Rate, and Vessel Diameter on Precipitation Formation

#### 3.2.1 Varying compound concentration

As a first step towards understanding the physical parameters affecting compound precipitation, we began by varying the concentration of ABT-737 from 0 to 50 mg/mL for a fixed 20 μL injection volume with Q_v_ = 4 mL/min, Q_i_ = 100 μL/min, and D_v_ = 2.9 mm. These values were chosen as an empirical starting point at which a substantial amount of precipitation was observed. A single escalating concentration experiment used a total of 110 μL of vehicle, 110 μL of ABT-737 at 50 mg/mL, and 71.61 mL of blood. A set of three replicate escalating concentration experiments were run in which the CPD was cleaned only after the final concentration. As a result, the panels in Figs. 5A, 5B, and 5C depict the cumulative precipitate trapped for all prior concentrations.

**Fig. 5.**
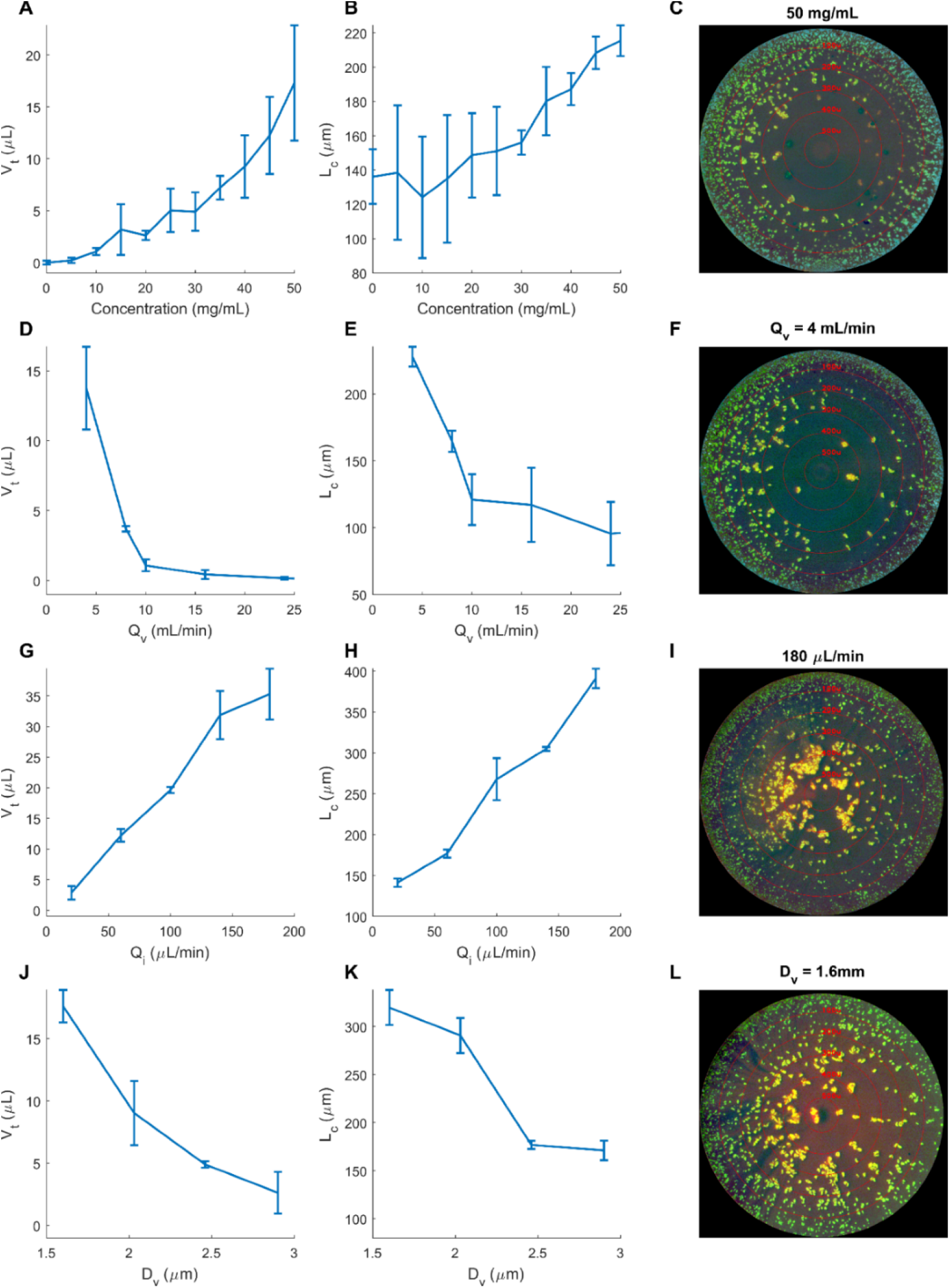
Study of the effect of compound concentration (A-C), vessel flow rate (D-F), compound infusion rate (G-I), and vessel diameter (J-L) on precipitation formation. Data points show the mean and standard deviation (n=3). In each case 20 μL of ABT-737 is injected into the IB. A) and B) show the effect of concentration on total particle volume and the characteristic size. C) shows a representative differential image of the CPD at 50 mg/mL. Compound is injected with Q_i_ = 100 μL/min into whole blood flowing at Q_v_ = 4 mL/min with D_v_ = 2.9 mm. D) and E) show the effect of vessel flow rate on total particle volume and the characteristic size. F) shows a representative differential image of the CPD at Q_v_ = 4 mL/min. Compound is injected at 50 mg/mL is injected with Q_i_ = 100 μL/min into whole blood flowing through an IB with D_v_ = 2.9 mm. G) and H) show the effect of injection flow rate on total particle volume and the characteristic size. I) shows a representative differential image of the CPD at Q_i_ = 180 μL/min. Compound is injected into whole blood with Q_v_ = 10 mL/min and D_v_ = 2.9 mm. J) and K) shows the effect of vessel diameter on total particle volume and the characteristic size. L) shows a representative differential image of the CPD for D_v_ = 1.6 mm. The flow rates are adjusted so that the average linear velocity is 75 cm/s and the ratio of Q_v_/Q_i_=50.

With increasing compound concentration, the amount and size of precipitate both increase (Fig. 5A and 5B). Starting at 0 mg/mL, only vehicle is introduced into the system and no precipitate can form. However, a small number of particles are quantified due to false positives detected by the CNN. We consider this a negligible noise floor that can improve with better cameras and further training data. With escalating compound concentration, *V*_*t*_ grows with a roughly power-law increase as the system becomes more supersaturated and more compound becomes available to precipitate out of solution. Fig. 5B shows a corresponding increase in *L*_*c*_ from 124 μm at 10 mg/mL to 215 μm at 50 mg/mL. At lower concentration there is more variability in *L*_*c*_ quantitation due fewer particles being present to provide a statistical sample. To acquire a more stable measurement at these lower concentrations, either more replicates can be performed, or a larger injection volume could be used, provided the CPD does not become saturated. A representative differential image of the CPD after the 50 mg/mL infusion is shown in Fig. 5C.

#### 3.2.2 Varying vessel flow rate

We next sought to isolate the effect of Q_v_ by infusing 20 μL of compound at 50 mg/mL concentration and 100 μL/min Q_i_ while varying Q_v_ (Figs. 5D, 5E, and 5F). 3 replicates were acquired for each point and the CPD was cleaned between each acquisition. A single set of 5 Q_v_ values used a total of 100 μL of ABT-737 at 50 mg/mL and 62.78 mL of blood.

For Q_v_ greater than 24 mL/min, the amount of compound delivered to the system per unit time is small compared to the amount of blood that flows past the injection point. As a result, no appreciable amount of precipitate is formed at these high flow rates. As Q_v_ decreases, the relative amount of compound near the injection site increases and precipitate forms at a sharply accelerating rate (Fig. 5D). For flow rates lower than 4 mL/min, the IB generated so much precipitate that the flow path became clogged, and no measurements could be acquired. Similarly, with decreasing Q_v_, *L*_*c*_ increases to an average value of 228 μm at 4 mL/min (Fig. 5E). A representative differential image of the CPD for Q_v_ = 4 mL/min is shown in Fig. 5F.

The increased production of precipitate at lower Q_v_ can, in part, be understood by considering the cross-sectional area that the compound and whole blood streams occupy downstream from the injection site. When the ratio of Q_i_ to Q_v_ is small, the cross-sectional area occupied by the compound stream is low. As a result, the compound can both quickly diffuse into blood and develop a low average concentration that is not conducive to precipitate formation. As the ratio of Q_i_ to Q_v_ increases, the compound stream occupies a larger cross-sectional area that takes a longer time to diffuse into the blood, allowing more chance for precipitate to form. Moreover, there is a larger reservoir of compound that generates a higher average concentration on which to form more and larger particles.

#### 3.2.3 Varying injection flow rate

We next studied the effect of varying the compound infusion rate. A fixed compound volume of 20 μL was injected at 50 mg/mL into whole blood flowing with Q_v_ = 4 mL/min while varying Q_i_. 3 replicates were acquired for each point and the CPD was cleaned between each acquisition. A single set of 5 Q_i_ values used a total of 100 μL of ABT-737 at 50 mg/mL and 44.35 mL of blood.

With higher Q_i_, a larger amount of compound is added to the blood per unit time, resulting in more precipitate volume (Fig. 5G) with larger size particles (Fig. 5H). This effect can also be explained as above by considering the relative cross-sectional area of the streams and average system concentration as the ratio of Q_i_ to Q_v_ increases. A representative differential image of the CPD for Q_i_ = 180 μL/min is shown in Fig. 5F.

Interestingly, *V*_*t*_ surpasses the 20 μL of solubilized compound that was injected into the system. This is likely due to an overestimate of the volume calculated by the image analysis of the CPD due to two main factors. First, when larger particles become trapped in the CPD they block the flow path for later smaller particles, potentially causing a subset of particles’ size to be overestimated. Second, compound precipitates are non-spherical and can have a very jagged shape. As a result, the treatment of a single particle’s volume as the summation of several rectangular prisms (as in Eq. 2-4) can be an overestimate.

#### 3.2.4 Varying vessel diameter

As a final experiment to explore the parameter space available for quantification by OSPREY, we fabricated 4 IB’s with D_v_ = 1.6, 2.04, 2.46, and 2.9 mm. To make measurements from the different D_v_ more comparable, we scaled Q_v_ such that the average linear velocity was 75 cm/s, and the ratio of Q_v_/Q_i_ was 50. Despite keeping these other parameters constant, both the amount (Fig. 5J) and size (Fig. 5K) of precipitate decreased as D_v_ increased. A representative differential image of the CPD for D_v_ = 1.6 mm is shown in Fig. 5L.

We attribute the increased precipitation for smaller D_v_ to the increased shear stresses acting on the fluid streams creating local nucleation points(Sakariassen et al., 2015). We note that although whole blood must be treated a non-Newtonian fluid for D_v_<0.3 mm, for the larger sizes in this experiment it can be considered as Newtonian(Ascolese et al., 2019). As a result, the viscosity of the whole blood is the same for all D_v_.

### 3.3 Demonstration of precipitation in two physiologically relevant model systems

As a final demonstration of the potential applicability of OSPREY to modeling *in-vivo* effects in an *in-vitro* system, we performed experiments using physiologically relevant estimates of Q_v_ and D_v_ that would be found in a rat femoral vein (Q_v_ = 2.5 mL/min and D_v_ = 1.6 mm) and dog saphenous vein (Q_v_ = 65 mL/min and 2.9 mm D_v_). The compound infusion rate was 75 μL/min in rat and 100 μL/min in dog. Fig. 6A shows the results for total particle volume as the compound concentration was progressively increased. Although no appreciable amount of precipitate was detected in the dog model even up to 40 mg/mL compound concentration, in the rat model precipitate formation began at 5 mg/mL and increased monotonically from there. Figs. 6B and 6C show the final differential images for the rat and dog models, respectively, at the 40 mg/mL concentration with insets showing an image of the IB during experiment acquisition. Qualitatively, while the CPD is filled with precipitate in the rat model, there is little to no precipitate apparent in the dog model. The rat model experiment was stopped before reaching the full 50 mg/mL concentration due to a saturation of the CPD with particles.

**Fig. 6.**
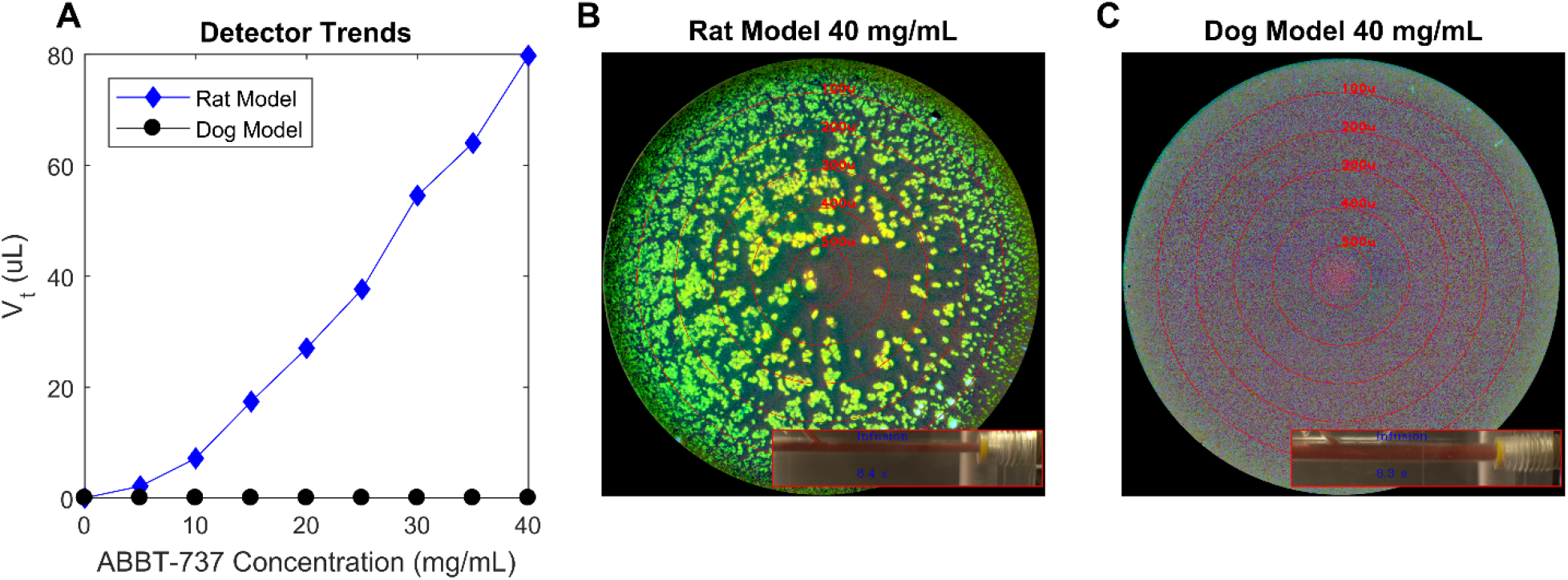
Modeling precipitation with physiologically relevant Qv and Dv for rat femoral vein and dog saphenous vein. ABT-737 was injected into whole blood with Qv = 2.5 mL/min and Qi = 75 μL/min for the rat model and Qv = 65 mL/min and Qi = 100 μL/min for the dog model. A) Total particle volume as a function of compound concentration in rat and dog models. A single replicate was acquired at each concentration. B) Differential image for the rat model system at 40 mg/mL compound concentration. Inset shows the IB with 1.6 mm model vessel diameter. C) Differential image for the dog model system at 40 mg/mL compound concentration. Inset shows the IB with 2.9 mm model vessel diameter.

The lack of precipitate in the dog model as compared with the plentiful precipitate in the rat model can be qualitatively understood by comparing the run parameters between the two experiment sets. While the increased Q_i_ (+33.3%) in the dog model typically supports a higher amount of precipitate generation, this is more than countered by the increase in Q_v_ (+2500%) and D_v_ (+81.25%) that tend to reduce precipitate formation. As a result, the dominating effects of high Q_v_ and D_v_ lead to essentially no precipitate in the dog model.

### 3.4 Discussion on the Application of OSPREY to Study Solubility Challenged Compounds Prior to IV Administration

Our primary rationale for developing OSPREY is to better understand the dynamic precipitation processes taking place during preclinical evaluation of novel compounds by IV infusion. By better understanding these effects and cataloguing their dependencies on various compound classes and formulations, we can better triage prospective compounds for further investigation. Moreover, OSPREY offers an *in-vitro* platform on which to assess and modify compound administration protocols for better *in-vivo* outcomes. Taken together, we envision the use of OSPREY as a method to pre-qualify solubility challenged compounds in preclinical cardiovascular and toxicology studies. In this way, OSPREY can contribute to the reduction principle of animal welfare by eliminating uninformative experiments due to precipitation events, avoid generation of discouraging safety data that is not pharmacologically relevant, allow test compound to be more judiciously used, and reserve *in-vivo* resources for informative experiments.

The demonstration in section 3.3 serves as an example of one way in which OSPREY can be used to assess compound formulation prior to preclinical IV infusion. In this workflow, an IB can be made to match the physiological diameter of an animal model blood vessel of interest and an escalating dose experiment can be run using the corresponding physiological flow rate. An appropriate *in-vivo* infusion concentration can then be determined by remaining below the concentration at which a substantial amount of precipitation is observed. Similar experiments can also be made to determine an appropriate infusion rate. Based on the results in this paper, very different amounts of precipitation can be expected when administering a compound in blood vessels with different size and flow rate. Therefore, even if a specific administration procedure is acceptable in one animal model system, it may not be in another.

OSPREY possesses several appealing aspects that make it a valuable research tool for quantifying dynamic precipitation events. These include the ability to quantify particle sizing in whole blood, the ability to rapidly change the 3D printed IB to model different systems, automated interrogation of various system parameters, and automated image analysis using a trained CNN model. At the same time, there are several caveats that should enumerated. First, the IV injection site modeled in the IB cannot fully recapitulate the complex fluidic path and flow profiles that occur *in-vivo*. For instance, blood flowing in the infused vein transitions into the larger vena cava and passes through turbulence in the heart before traveling to the pulmonary arteries where precipitate accumulates *in-vivo*. Future modifications to the IB can build more complexity into the existing OSPREY system but will nevertheless be an imperfect version of the *in-vivo* environment. The next caveat is inaccuracies in quantitation that result from measuring non-spherical and jaggedly shaped precipitates in the CPD. Accurate quantification of *V*_*t*_ and *L*_*c*_ in the CPD relies on particles becoming wedged flat against the viewing window. However, actual precipitates may only contact the viewing window at one point – so their actual size may differ from where they become trapped. Moreover, with high enough Q_v_, a trapped particle can be forced to change shape and move to a new location. A third caveat is that the acquired data is highly dependent on the quality of the whole blood and compound solution introduced into the device. We have seen variability in results depending on how the whole blood is processed and stored before experiment. Similarly, we have seen some presumably thermodynamically metastable compound solutions precipitate in static conditions over the course of a day before being analyzed on OSPREY.

## Conclusions

This report describes the development and characterization of OSPREY, a new flow-through imaging device for modeling precipitation of prospective small molecule drugs in an *in-vitro* system. The automated experiment acquisition and image processing enabled us to study the effects of compound concentration, Q_v_, Qi, and D_v_ on the formation of drug precipitates. We further provided a demonstration of the ability of OSPREY to model *in-vivo* systems using physiologically relevant system parameters for modeling rat femoral vein and dog saphenous vein.

We view this first report on OSPREY as a steppingstone towards realizing its potential to prequalify solubility challenged compounds for *in-vivo* administration. To reach this goal, further studies into parameters related to the compound formulation (e.g., pH, additives, viscosity, density, etc.) and compound classes (e.g., beyond rule-of-5, amorphous, crystalline, etc.) found in drug development pipelines are needed, as well as the correlation between these results and *in-vivo* experiments.

## Supporting Information

Additional experimental details including photographs and videos of experimental setup.

## Supporting information

CPD blood video

CPD plasma video

Injection block blood video

Injection block plasma video

## Abbreviations

OSPREY: optical spatial precipitation analyzer
CPD: conical particle detector
IB: injection block
CNN: convolutional neural network
IV: intravenous
Q_v_: model vessel flow rate
Q_i_: compound infusion flow rate
D_v_: model blood vessel diameter
V_t_: total particle volume
L_c_: characteristic particle size

## Acknowledgements

The authors acknowledge Drew Wollman of AbbVie for his helpful discussions on fluid flow and improving instrument design. The work was enabled by the AbbVie Experiential Intern Program. AJR, RLM, WRB, AW, and JYP are employees of AbbVie. LH was an employee of AbbVie at the time of the study. The design, study conduct, and financial support for this research were provided by AbbVie. AbbVie participated in the interpretation of data, review, and approval of the publication.

