## Supplemental Information for "In-vitro Modeling of Intravenous Drug Precipitation by the Optical Spatial Precipitation Analyzer (OSPREY)"

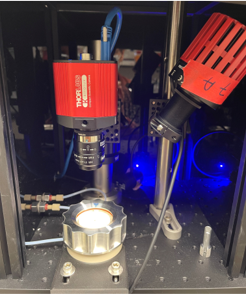

Fig S1 Close up of CPD, LED illumination, and camera

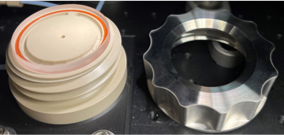

Fig S2 CPD deconstructed

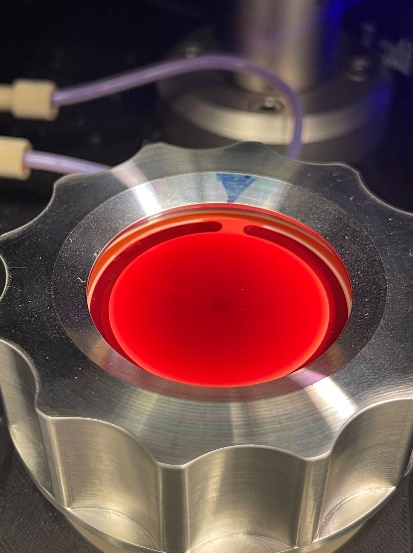

Fig S3 CPD with whole blood

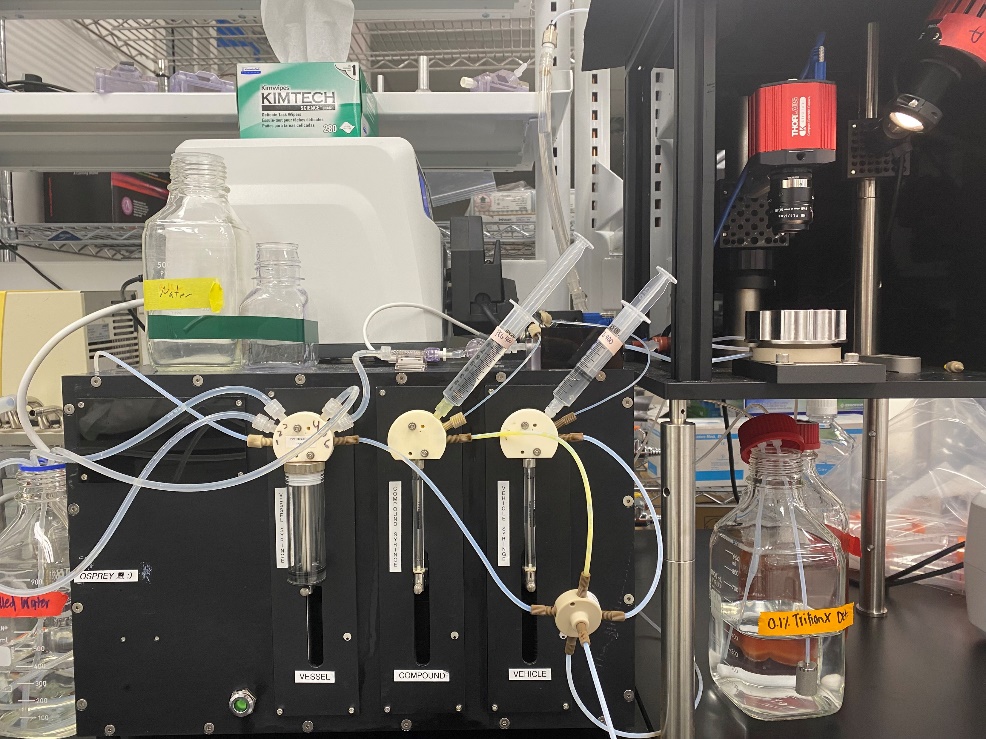

Fig S4 Image of the 3-syringe system

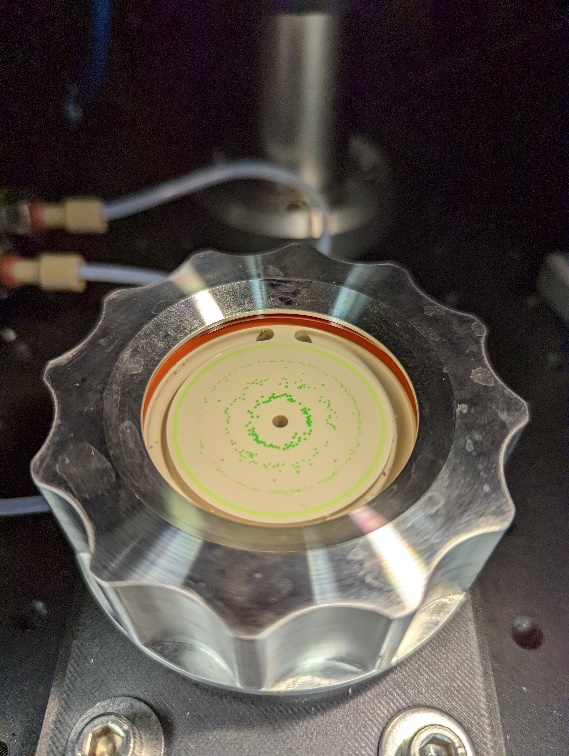

Fig S5 Image of the CPD with calibration microspheres

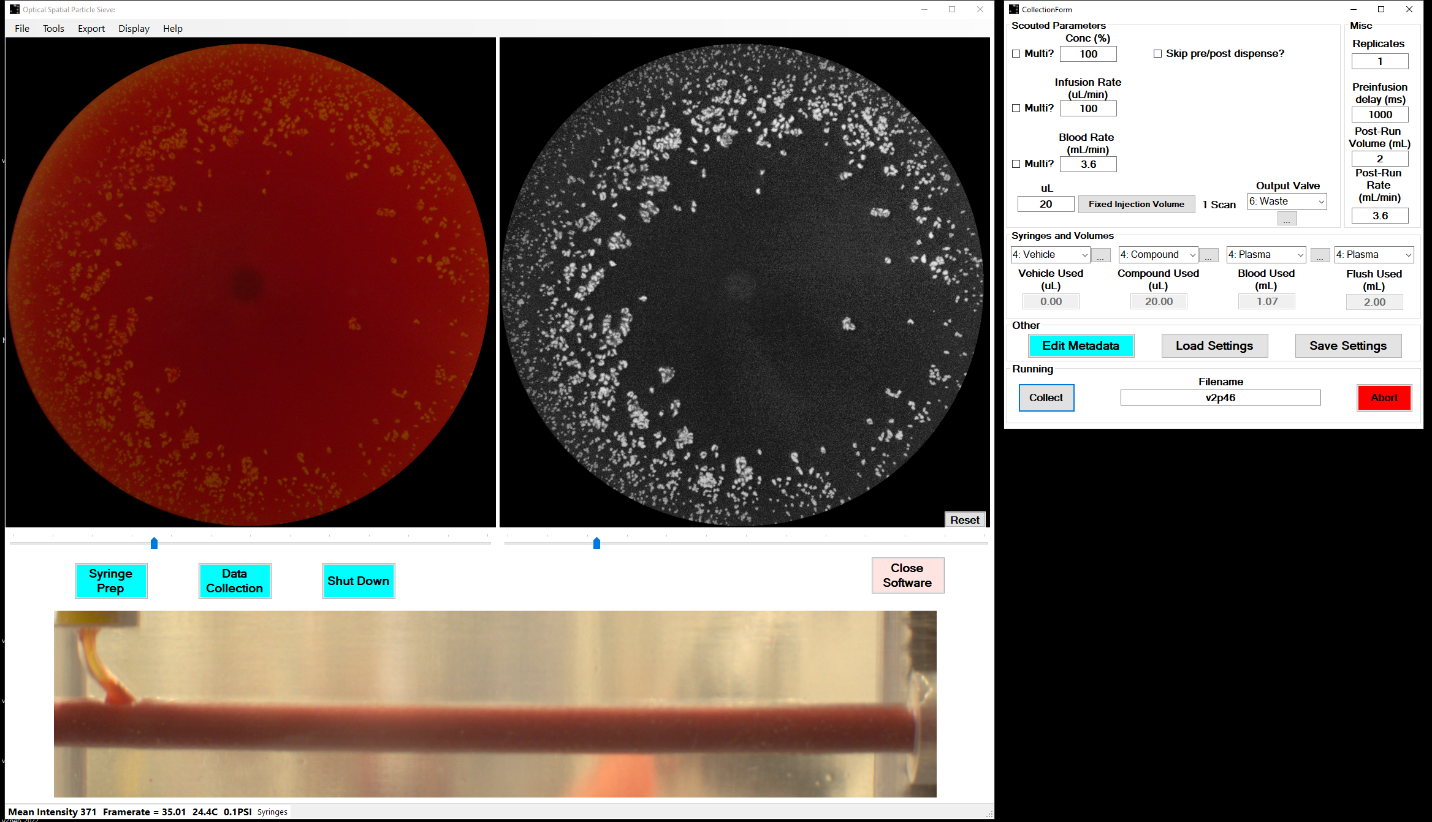

Fig S6 Screenshot of OSPREY acquisition software during experiment

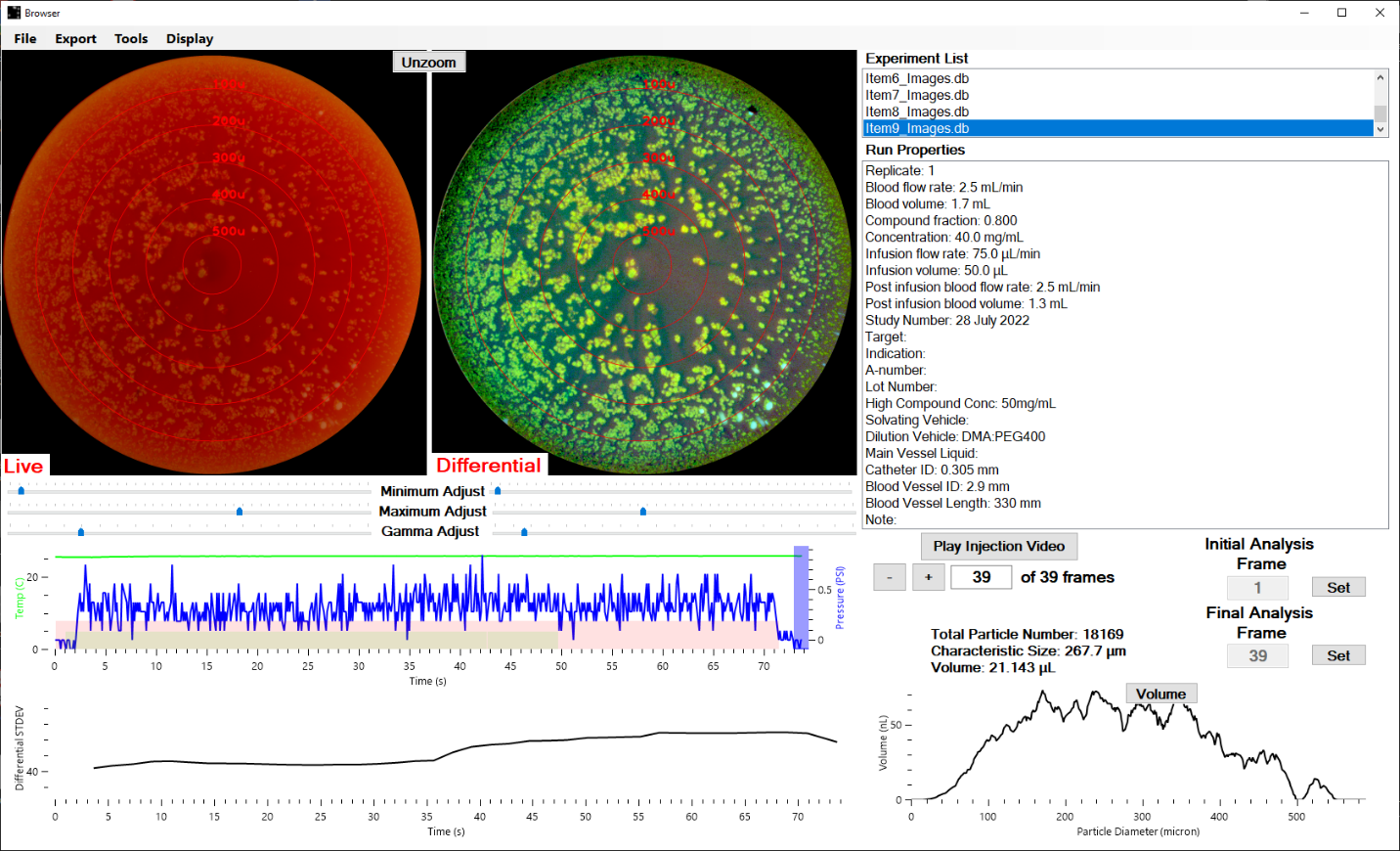

Fig S7 Screenshot of OSPREY browser software

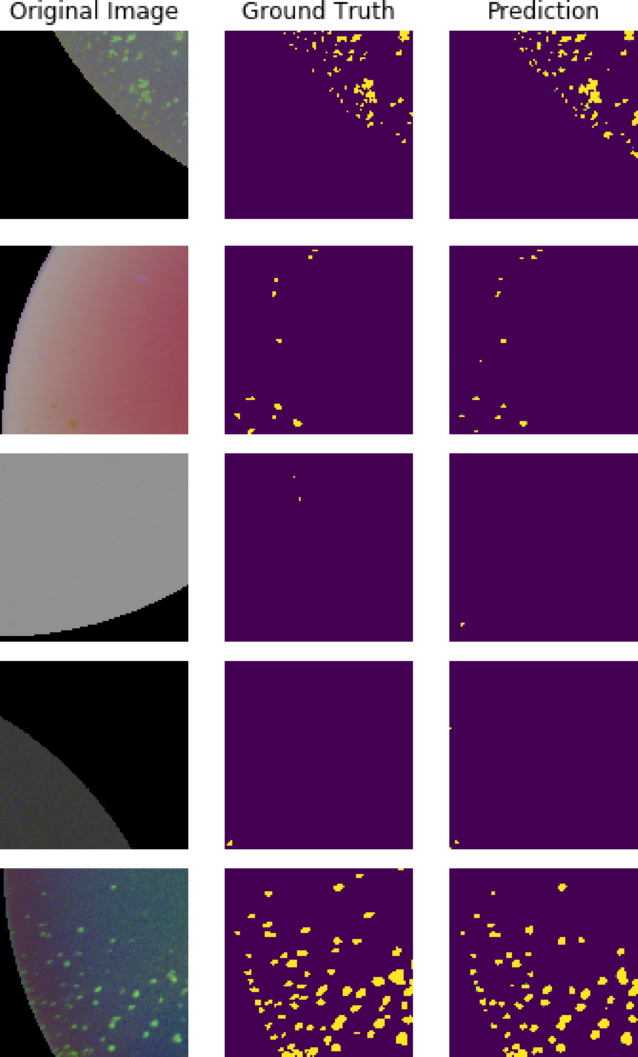

Fig S8 Original, ground truth, and predicted mask images from a reserved validation set. (Images split across two pages. Part 1 of 2 in Fig S8)

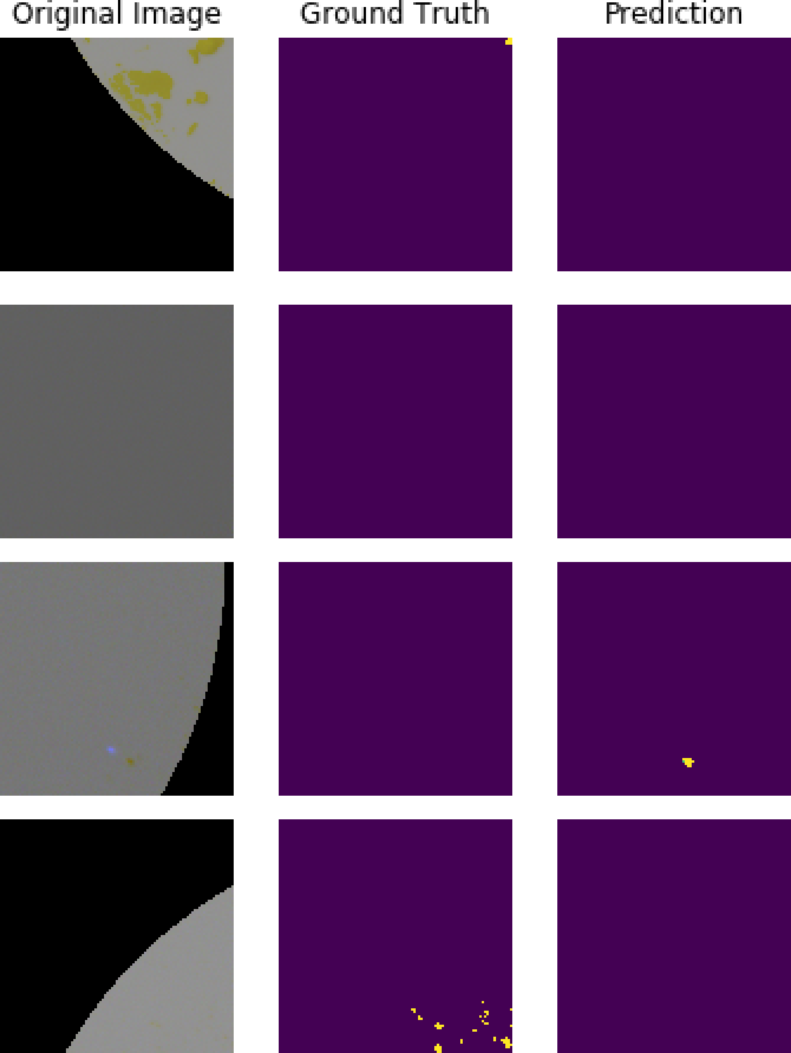

Fig S9 Original, ground truth, and predicted mask images from a reserved validation set. (Images split across two pages. Part 2 of 2 in Fig S9)
